# Cryo-electron microscopy structure of the H3-H4 octasome without histones H2A and H2B

**DOI:** 10.1101/2021.10.27.466091

**Authors:** Kayo Nozawa, Yoshimasa Takizawa, Leonidas Pierrakeas, Kazumi Saikusa, Satoko Akashi, Ed Luk, Hitoshi Kurumizaka

## Abstract

The canonical nucleosome, which represents the predominant packaging unit in eukaryotic chromatin, has an octameric core made up of two histone H2A-H2B and H3-H4 dimers with ~147 base-pair (bp) DNA wrapping around it. Non-nucleosome particles with alterative histone stoichiometries and DNA wrapping configurations have been found, and they could profoundly influence genome architecture and function. Here we solved the structure of the H3-H4 octasome, which is a nucleosome-like particle with a core made up of four H3-H4 dimers. Two conformations, open and closed, are determined at 3.9 Å and 3.6 Å resolutions by cryo-electron microscopy, respectively. The H3-H4 octasome, made up of a di-tetrameric core, is wrapped by ~120 bp DNA in 1.5 negative superhelical turns. The symmetrical halves are connected by a unique H4-H4’ interface along the dyad axis. *In vivo* crosslinking of cysteine probes placed at another unique H3-H3’ interface demonstrated the existence of the H3-H4 octasome in cells.

## Introduction

Eukaryotic genomic DNA is packaged into chromatin, in which the basic unit is the nucleosome core particle, consisting of two histone H2A-H2B and H3-H4 dimers forming a histone octamer with a 145-147 base-pair (bp) DNA fragment wrapped around in 1.7 left-handed superhelical turns (Luger et al., 1997). The formation of the nucleosome limits the accessibility of the underlying DNA sequence and is therefore generally inhibitory to the binding of sequence specific factors (Kornberg and Lorch, 1999). Nucleosomes are structurally heterogeneous inside the cell. The core histones can be replaced by histone variants or decorated with covalent modifications, generating a repertoire of structurally distinct nucleosomes along the chromatin with altered biophysical and biochemical properties that contribute to the regulation of chromosome structure and nuclear activities (Jenuwein, 2001; Kurumizaka et al., 2021).

While the nucleosome represents the predominant histone-DNA assembly in cellular chromatin, the existence of histone-DNA complexes with alternative stoichiometries, such as sub-nucleosomes, has been implicated (Lai and Pugh, 2017; Luger et al., 2012). For example, the hexasome, which has one less H2A-H2B dimer than the canonical nucleosome, can form when RNA polymerase II (RNAPII) transits through a nucleosome, displacing one of its two H2A-H2B dimers (Arimura et al., 2012; Ramachandran et al., 2015). Since RNAPII passage on the nucleosomal DNA itself does not release the H2A-H2B dimer (Ehara et al., 2019; Kujirai et al., 2018), the transcription-mediated H2A-H2B removal may require additional activity such as histone chaperones, nucleosome remodelers, histone variants, and/or histone modifications (Venkatesh and Workman, 2015). The tetrasome, which contains an (H3-H4)_2_ tetramer core without H2A-H2B, is an intermediate structure involved in nucleosome formation (Jorcano and Ruiz-Carrillo, 1979). During nucleosome formation, two H3-H4 dimers associate with a DNA fragment to form a tetrasome followed by the deposition of two H2A-H2B dimers in a process that is mediated by histone chaperones *in vivo* (Böhm et al., 2011; Mattiroli et al., 2017b).

The H3-H4 tetrasome, which is associated with ~70 bp DNA, can be reconstituted by mixing (H3-H4)_2_ tetramers and DNA at equimolar ratio (Dong and van Holde, 1991; Lavelle and Prunell, 2007). At higher protein-to-DNA ratios, H3-H4 can form a nucleosome-size particle consisting of an octameric H3-H4 core wrapped around by ~130 bp of DNA (Moss et al., 1977; Simon et al., 1978). This nucleosome-like particle, called the H3-H4 octasome hereafter, exhibits a bead-like structure with a diameter comparable to the nucleosome, as evidenced by early electron microscopy (EM) studies and a recent atomic force microscopy analysis (Bina-Stein, 1978; Lavelle and Prunell, 2007; Zou et al., 2018). The (H3-H4)_2_ tetramer alone is sufficient to position around the center of the *Lytechinus variegatus* 5S rDNA sequence, a naturally occurring nucleosome positioning sequence (Dong and van Holde, 1991). Interestingly, reconstitution under a 2:1 tetramers-to-DNA concentration, the tetramers re-distribute equally to the two halves of the 5S rDNA sequence to form a ‘di-tetrasome’ particle, consistent with the H3-H4 octasome configuration (Flaus et al., 1996).

In this study, we determined the cryo-EM structures of the H3-H4 octasome. The data revealed that the core of H3-H4 octasome is made up of two (H3-H4)2 tetramers, which are wrapped around by ~120 bp DNA in 1.5 left-handed superhelical turns. Along the dyad axis where the two tetramers meet, two H4 molecules form a unique four-helix bundle. To assess the biological relevance of the H3-H4 octasome structure, we interrogated yeast chromatin with an *in vivo* disulfide crosslinking assay and detected an H3-H4 octamer-specific interaction between two inward facing H3 molecules at the disk-disk interface. The implications of the H3-H4 octasome on chromatin architecture and genome function are discussed.

## Results

### Reconstitution of the H3-H4 octasome

We first reconstituted the nucleoprotein complexes with the human histones H3 and H4 and a 145 bp DNA fragment containing the Widom 601 positioning sequence by the salt dialysis method. Two types of complexes were reconstituted and then separated by native polyacrylamide gel electrophoresis (PAGE) (Figure 1A). Electrospray ionization mass spectrometry revealed that the molecular weights of the nucleoproteins corresponding to the upper and lower bands are 149,789 (±48) and 201,681 (±71), which are consistent with the theoretical molecular weights for the H3-H4 tetrasome and the H3-H4 octasome, respectively (Figure 1B-C).

**Figure 1.**
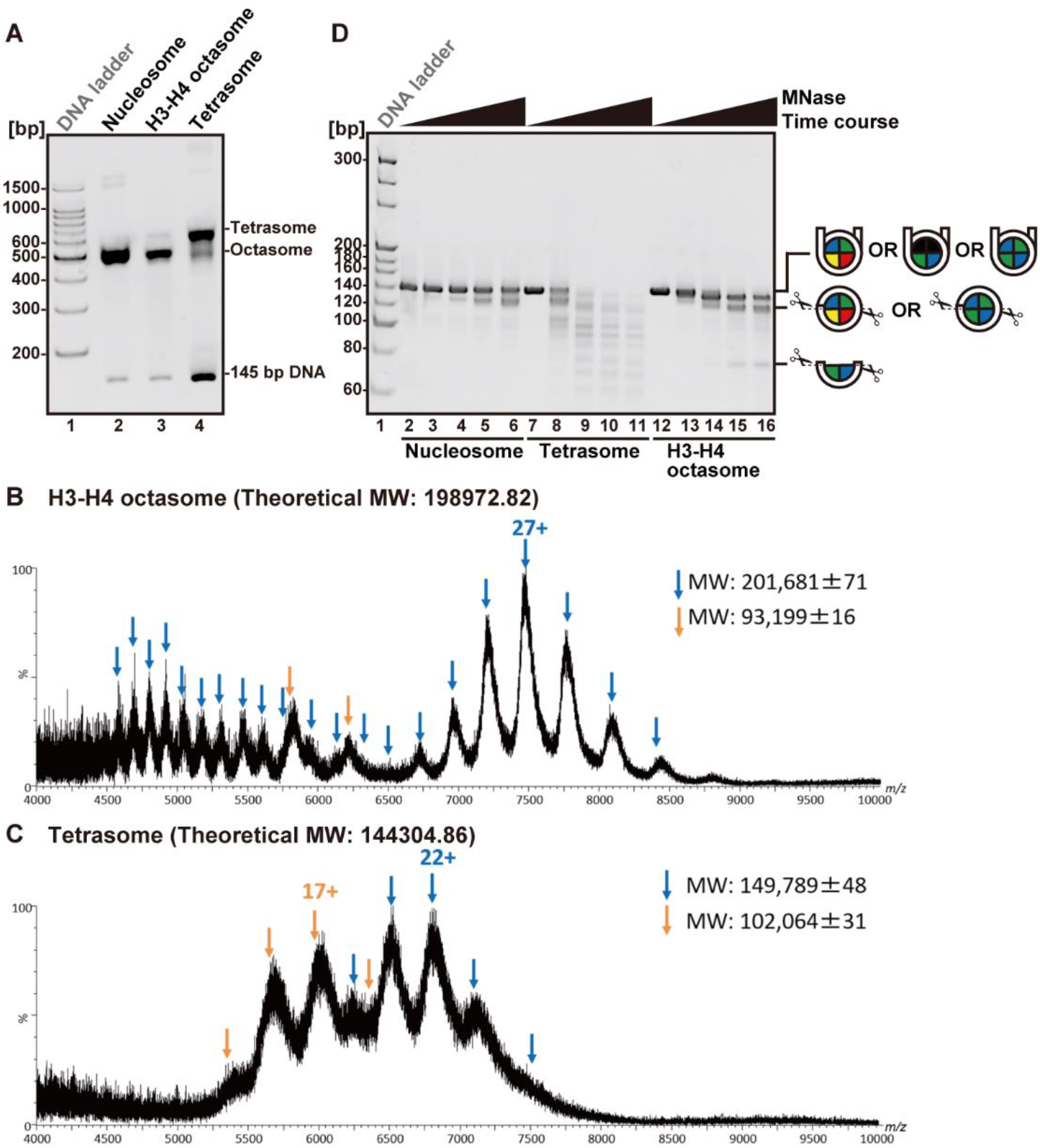
Reconstitution of the H3-H4 octasome. (A) Purified nucleosomes, H3-H4 octasomes, and tetrasomes (200 ng each) reconstituted with human histones and a 145-bp Widom 601 positioning sequence were analyzed by native PAGE and ethidium bromide (EtBr) staining. ESI mass spectra of H3-H4 octasomes (B) and tetrasomes (C). Blue and orange arrows indicate multiply charged ions of the nucleosomes and double stranded DNA, respectively. Numeric values indicate the charge state of the dominant peak for individual species. (D) The indicated nucleoproteins were treated with MNase for 0, 3, 6, 18, and 36 min from left to right. The DNA products were dissociated from the histones and analyzed by native PAGE with EtBr staining.

To assess the extent of histone-DNA contacts within the H3-H4 octasome, we performed a nuclease sensitivity assay on H3-H4 octasomes, along with canonical nucleosomes and H3-H4 tetrasomes as controls (Figure 1D). Micrococcal nuclease (MNase) is an endo/exo nuclease that preferentially digests DNA detached from the histone surface, but not DNA stably wrapped around histones, as in the nucleosome (Noll, 1974). Time courses of MNase digestion show that the canonical nucleosome protects a dominant DNA species at ~145 bp, corresponding to the stably wrapped nucleosomal DNA, and a ~120-bp species at later time points, consistent with partial unwrapping of the DNA ends (Koopmans et al., 2009) (Figure 1D, lanes 2-6). By contrast, the H3-H4 tetrasome was highly susceptible to MNase attack, suggesting substantial exposure of its DNA to solvent (Figure 1D, lanes 7-11). The H3-H4 octasome exhibited a much stronger protection of the DNA compared to the H3-H4 tetrasome, although the full-length 145 bp DNA fragment was progressively trimmed to ~130 bp and further to ~120 bp and 70 bp at later time points (Figure 1D, lanes 12-16). The DNA protection pattern of the H3-H4 octasome suggests that the octasomal histone core is stably wrapped by a DNA fragment but in a manner that differs from the canonical nucleosome.

### Overall structure of the H3-H4 octasome

We determined the structure of the H3-H4 octasome by cryo-electron microscopy (Figure 2A, figure supplement 1A-C). The purified H3-H4 octasome sample was subjected to data collection with a 300 kV electron microscope, and 1.43 million particles related to the H3-H4 octasome were identified from 5,517 electron micrographs. Two distinct H3-H4 octasome structures, closed and open forms, were determined by single particle analyses at 3.6 Å and 3.9 Å resolutions, respectively (Figure 2B, figure supplement 1D). In both the closed and open forms, a 120 bp DNA segment wraps around the octameric H3-H4 core 1.5 times in a left-handed manner, resembling a clamshell. The opening measured between the two DNA gyres at the farthest points from the H3-H4 octasomal dyad is 7.7 Å wider in the open form compared to the closed form (Figure 2C). In both structures, two H3-H4 tetramers are connected by a unique H4-H4’ four-helix bundle (FHB) that coincides with the octasomal dyad. Each H3-H4 tetramer constitutes a disk that represents a symmetrical half of the H3-H4 octasome. In the nucleosome, the FHB at the H3-H3’ interface, which links the two H3-H4 dimers, is located at the nucleosomal dyad; however, in the H3-H4 octasome, there are two H3-H3’ FHBs, and they are positioned three helical turns away from the H3-H4 octasomal dyad at the superhelical locations (SHLs) +3 and −3 (Figure 2D). These histone relocations enable each H3-H4 tetramer to symmetrically engage a ~60 bp DNA segment on either side of the dyad (SHL0) in the H3-H4 octasome.

**Figure 2.**
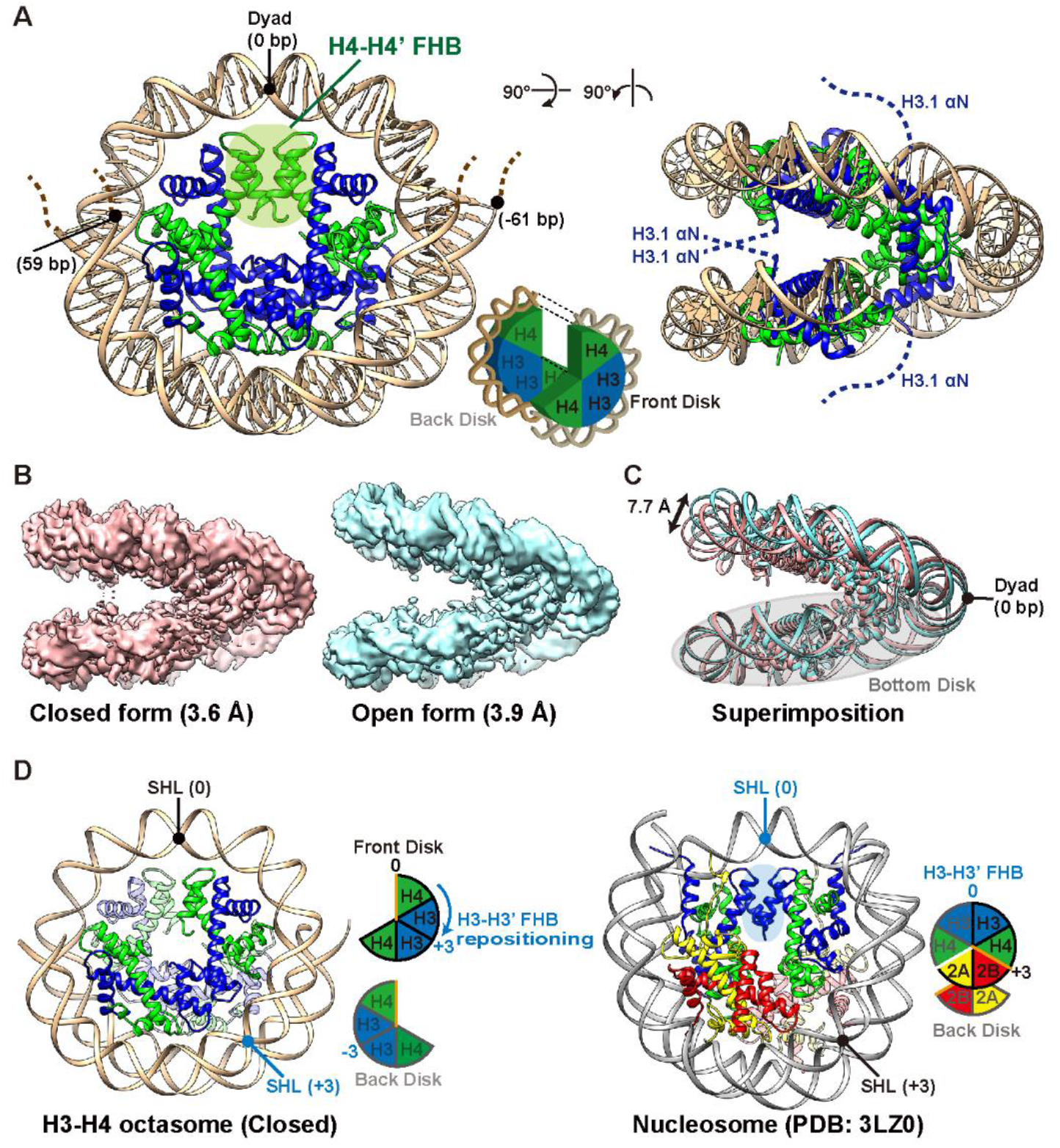
Cryo-EM structures of the H3-H4 octasome. (A) The cryo-EM structure of the H3-H4 octasome (closed form). The electron densities of the N-terminal regions (amino acid residues 1-58) of H3.1 and the DNA termini were not observed (indicated by dashed lines). (B) Cryo-EM densities of the H3-H4 octasome in the closed (left panel) and open (right panel) forms. (C) Superimposition of the open and closed forms of the H3-H4 octasome, performed on the bottom disk. The double arrow indicates the difference of the measured distances between the DNA gyres in the open and closed forms. (D) Ribbon models of the H3-H4 octasome (left panel) and the canonical nucleosome (right panel). In the H3-H4 octasome, the two H3-H3’ FHBs are positioned three helical turns away from the H3-H4 octasomal dyad at the superhelical locations (SHLs) +3 and −3. Orange lines in the schematic representations of the histone core assemblies indicate the H4-H4’ interface and the H4-H2B interface of the H3-H4 octasome and the nucleosome, respectively. The white line between H4 and H2A in the nucleosome schematics emphasizes the two proteins are not connected.

Structural comparison of the H3-H4 octasome with the canonical nucleosome revealed additional H3-H4 octasome-specific features. First, the H3-H4 octasome lacks the acidic patch provided by H2A and H2B, a docking site for a variety of nucleosome binding proteins (Sundaram and Vasudevan, 2020) (Figure 3A). Second, the opening between the two disks of the H3-H4 octasome, even in the closed form, is wider than that of the nucleosome, leading to a ~20 Å extension between the DNA gyres of the H3-H4 octasome, as compared to the nucleosome (Figure 3B, right panel). Third, the αN region of H3 is predicted to occupy the inter-disk space of the H3-H4 octasome; however, the expected alpha helical structure is not visible, suggesting that the H3 αN regions in the inter-disk space are dynamic. The remaining two H3 αN regions exposed on outer surfaces of the H3-H4 octasome are also unstructured. Finally, the FHB at the octasomal H4-H4’ interface resembles the one at the nucleosomal H4-H2B interface (figure supplement 2A-C). For example, in the nucleosome, the sidechains of H4 Tyr72, Glu74, His75, and Arg92 interact with those of H2B Glu76, Arg99, Glu93, and Leu100, respectively. In the H3-H4 octasome, the same H4 sidechains are oriented in a similar manner, but they interact with the sidechains of Arg92, Tyr88, Asp85, and Asp68 on the opposite H4’ instead (Figure 3C).

**Figure 3.**
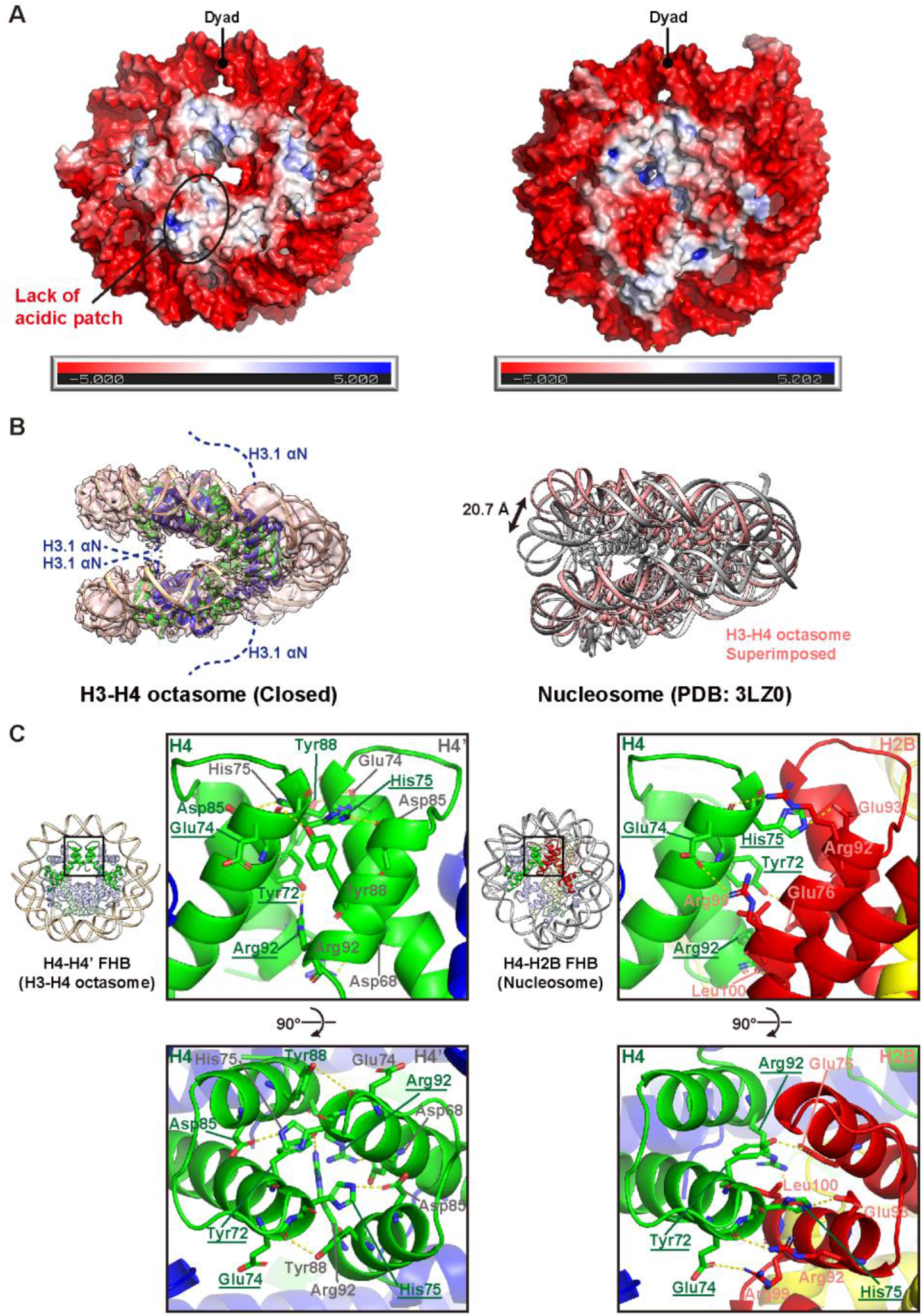
Structural comparison of the H3-H4 octasome and nucleosome. (A) Surface electrostatic potentials of the H3-H4 octasome (left panel) and the nucleosome (right panel). (B) The cryo-EM density map and the ribbon model of the H3-H4 octasome (closed form) are superimposed (left panel). The ribbon model of the H3-H4 octasome and that of the nucleosome are superimposed (right panel). (C) Close up views of the H4-H4’ interface of the H3-H4 octasome (left panel) and the H4-H2B interface of the nucleosome (right panel). Common residues utilized in both interactions are underlined.

### Detection of H3-H4 octasome-specific interactions in yeast cells

To search for the existence of native H3-H4 octasomes, we performed *in vivo* disulfide crosslinking experiments in *Saccharomyces cerevisiae* based on the VivosX technique (Mohan et al., 2018). We substituted Arg49 of yeast H3 with a cysteine (R49C) such that when the inward facing H3 molecules on the two disks of the H3-H4 octasome come together, a cystine linkage can be formed (Figure 4A). Given the proximity (with a theoretical Cβ-Cβ’ distance of 12.8 Å) and dynamic nature of the Arg49 sites, inter-disk H3-H3’ crosslinking is expected for R49C H3-H4 octasomes, but not for R49C nucleosomes, as the nucleosomal Arg49 residues are separated by a distance that is prohibitive to disulfide crosslinking (Cβ-Cβ’ 54.3 Å) (Figure 4B).

**Figure 4.**
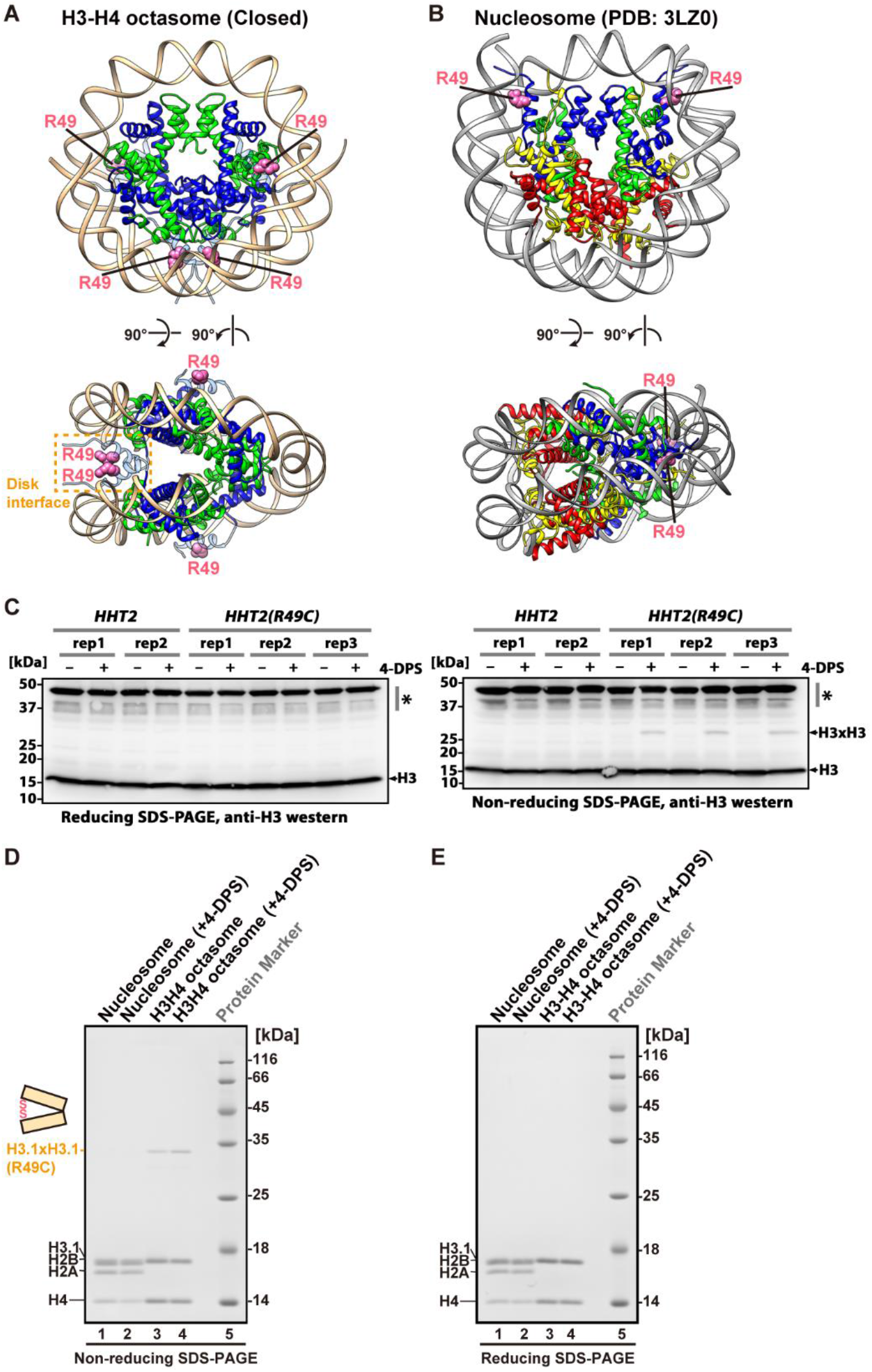
VivosX captures an interaction specific to the H3-H4 octasome in yeast cells. The locations of the H3 Arg49 residues in the H3-H4 octasome (A) and the nucleosome (B) are shown as pink spheres. The unobserved H3.1 αN helices of the H3-H4 octasome (pale blue) were modeled by superimposition with the H3.1 structure of the nucleosome (PDB ID: 3LZ0). (C) VivosX analysis of the H3 R49C yeast mutant and the wild-type H3 control. Protein extracts from cells treated with 4-DPS were subjected to reducing (left panel) and non-reducing (right panel) SDS-PAGE and analyzed by western blotting with an anti-H3 antibody. Non-specific bands are marked with asterisks. Biological replicates (rep) represent extracts prepared from individual transformants. Reconstituted nucleosomes and H3-H4 octasomes containing human histones and the C96S, C110A, and R49C substitutions on H3.1 were treated with or without 4-DPS and then subjected to non-reducing (D) and reducing (E) SDS-PAGE analyses with Coomassie Brilliant Blue staining.

To maximize the detection of any potential H3-H3’ crosslinking, R49C was introduced into the yeast <u>h</u>istone <u>H3</u> gene *HHT2* (which is transcribed as a divergent unit with the <u>h</u>istone <u>H4</u> gene *HHF2*) on a low-copy *TRP1* plasmid in a strain deleted for the endogenous H3-H4 genes (*HHT1-HHF1* and *HHT2-HHF2*). Since the strain was initially supplemented with the *HHT1-HHF1* gene on a *URA3* plasmid, 5-FOA treatment was used to remove the *URA3* plasmid to generate cells expressing *HHT2(R49C)* as the sole source (figure supplement 3A). The *HHT2(R49C)* cells were viable, albeit slow growing, indicating that the *HHT2(R49C)* allele is at least partially functional (figure supplement 3B). After treating *HHT2(R49C)* and the wild-type *HHT2* cells with 4,4’-dipyridyl disulfide (4-DPS), a cell-permeable oxidant, total proteins were separated by non-reducing SDS-PAGE and analyzed by anti-H3 western blotting. A ~30-kD band was observed in the *HHT2(R49C)* mutant, but not *HHT2* (Figure 4C, *left panel*). The 30-kD band was removed if the extracts were pretreated with beta-mercaptoethanol (bME) before electrophoresis (Figure 4C, *right panel*). These data suggest that the 30-kD adduct is formed between two H3 molecules (15 kD) with the R49C site via a cystine linkage.

One trivial explanation for the source of the 30-kD adduct is that the H3 R49C protein crosslinks with a 15-kD non-histone protein. To verify that the 30-kD band is indeed an adduct between two H3 molecules, we co-transformed a V5-tagged *HHT1(R49C)* plasmid with the untagged *HHT2(R49C)* plasmid into the histone knockout strain, generating a ‘hybrid’ mutant. With one or two V5 tags, the mobility of the heterotypic H3x^V5^H3 and homotypic ^V5^H3x^V5^H3 adducts is expected to increase stepwise relative to the untagged H3xH3 adduct. The non-hybrid *V5-HHT1(R49C)* alone strain gave a 37-kD band, marking the ^V5^H3x^V5^H3 adduct (figure supplement 3C, lane 1). The *HHT2(R49C)* alone strain gave the original 30-kD band, marking the untagged H3xH3 adduct (figure supplement 3C, lane 4). The hybrid strain gave a new band above and below the 30-and 37-kD marks that was observed by both anti-H3 and anti-V5 probing (figure supplement 3C, lanes 2 and 3, indicated by H3x^V5^H3). This result confirmed that the 30-kD band observed in the untagged R49C mutant is a disulfide adduct formed with two H3 molecules.

In theory, the H3 R49C site on a nucleosome (Figure 4B) could crosslink to a proximal R49C site on a neighboring nucleosome if the entry/exit DNA segments on the two nucleosomes become transiently unraveled (Armeev et al., 2021). To assess how much of the crosslinking in H3 R49C was due to inter-nucleosomal interactions, we reconstituted nucleosomes and H3-H4 octasomes bearing the R49C substitution in histone H3.1 and subjected the purified nucleoproteins to 4-DPS-induced crosslinking *in vitro* (figure supplement 3D-E). Since human H3.1 contains two naturally occurring cysteines (Cys96 and Cys110), these residues were replaced with a serine and an alanine, respectively, to avoid spurious crosslinking to R49C. The non-reducing SDS-PAGE analysis showed no detectable crosslinking with the R49C nucleosome but a ~30-kD band was observed for the R49C H3-H4 octasome (Figure 4D). Although a lesser amount of the 30-kD adduct was also observed in the absence of 4-DPS presumably caused by atmospheric oxidation, both 30-kD bands in the H3-H4 octasome lanes disappeared when the nucleoproteins were treated with bME (Figure 4E). This *in vitro* crosslinking experiment showed that the H3-H3’ disulfide adduct is specific to the H3-H4 octasome structure, further validating that the H3-H3’ crosslinking observed in the H3 R49C yeast is due to the formation of H3-H4 octasome or octasome-like structures.

## Discussion

Nucleosomes act as a physical barrier for DNA binding proteins, which regulate genomic DNA functions such as transcription, replication, recombination, and repair. In these processes, nucleosomes must be disassembled and reassembled. Dysregulation of these cycles leads to various diseases, including cancer (Burgess and Zhang, 2013). Subnucleosomes and nucleosome-like particles are dynamic structures that emerge and/or function during these cycles (Boeger et al., 2004; Rhee et al., 2014). In the present study, we determined the cryo-EM structures of a unique nucleosome-like particle, the H3-H4 octasome, which has unique features not found in the canonical nucleosome. In the structure, an approximately 120 bp DNA segment and four of the H3-H4 dimers form a stable core particle without H2A and H2B.

The H3-H4 octasome surface lacks the common foothold known as the acidic patch, which functions as an anchoring site for nucleosome binding proteins, including histone modifiers and nucleosome remodelers (Sundaram and Vasudevan, 2020). Thus, the presence of H3-H4 octasomes could interfere with the propagation of epigenetic marks and the spacing of nucleosomal arrays. In addition, the acidic patch of a nucleosome also serves as the docking site for the H4 N-terminal tail of a neighboring nucleosome, an interaction critical for chromatin compaction (Kalashnikova et al., 2013; Luger et al., 1997). The absence of the acidic patch in the H3-H4 octasome therefore could interrupt chromatin fiber formation. In addition, the unstructured N-termini on the two outer H3 molecules are expected to occupy the space above and below the H3-H4 octasome disks, interfering with stacking interactions between nucleosomes. Thus, chromatin fibers punctuated with H3-H4 octasomes will likely contribute to alternative higher-order conformations that could influence a wide variety of genomic functions.

The H3-H4 octasome exhibits a larger inter-disk spacing between DNA superhelical gyres compared to the nucleosome. This is likely contributed by the lack of H2A-mediated L1-L1’ interaction and the bolstering effect of the extended unstructured N-termini on the two inward facing H3 molecules. The inter-disk spacing appears dynamic, as evidenced by the closed and open conformations of the H3-H4 octasome structure. The wider inter-disk space and the dynamic nature of the clamshell structure could mean that the H3-H4 octasome will provide greater access to DNA binding factors, such as pioneer factors, that prefer to bind to the DNA along the gyre (Dodonova et al., 2020).

What give rise to the H3-H4 octasome is currently unknown. One mechanism may involve histone deposition pathways that utilize H3/H4-specific chaperones, such as Asf1, HIRA, and CAF-1, but exclude chaperones with H2A/H2B specificity, such as NAP1 and FACT (De Koning et al., 2007). Alternatively, H3-H4 octasome-like structures could form when a chromatin remodeler, like SWI/SNF, forces adjacent nucleosomes to overlap. Structural and biochemical studies have shown that at the interface of the two overlapping nucleosomes, an H2A-H2B dimer dissociates to generate a hexasome (Dechassa et al., 2010; Kato et al., 2017). And if the remodeler continues to push the hexasome towards the nucleosome, it is conceivable that the inward facing H2A-H2B dimer of the nucleosome could also dissociate, generating a second hexasome. As such, the two interacting hexasomes could form an interface that is reminiscent of the H3-H4 octasome. Although this configuration will still have two H2A-H2B dimers on the terminal sides of the H3-H4 octasome structure, our structural data and *in vivo* crosslinking data do not rule out such a possibility.

The H4-H4’ four-helix bundle is a structural feature unique to the H3-H4 octasome that is not seen in the nucleosome or other sub-nucleosome structures. However, we cannot detect the H4-H4’ interaction *in vivo* by crosslinking (data not shown). This is likely due to a limitation of the VivosX methodology in that efficient crosslinking of two proximal cysteine residues requires structural flexibility typically associated with loops, whereas disulfide crosslinking of α-helices, like those in the FHB, is highly inefficient (Thornton, 1981). Notwithstanding this technical caveat, our structural data have revealed a striking similarity between the FHBs at the H4-H4’ interface (in the H3-H4 octasome) and the H4-H2B interface (in the canonical nucleosome), underscoring the alternative role the H4 residues at the FHB play when H4 interacts with another H3-H4 dimer, instead of an H2A-H2B dimer. Indeed, the structural symmetry required for the formation of the H4-H4’ FHB was not appreciated until now.

Each of the two outward facing H4 molecules in the H3-H4 octasome has an unpaired half of a FHB (i.e. an α2-L2-α3 domain) that can in theory pair with another H4 on an H3-H4 dimer or tetramer, raising the possibility that H3-H4 dimers can polymerize along the DNA. The structure of such a hypothetical (H3-H4)-exclusive chromatin array could resemble that of the archaeal histone-DNA complex, in which DNA wraps around a polymer of archaeal histone homodimers to form a quasi-continuous superhelical structure (Bowerman et al., 2021; Mattiroli et al., 2017a; Sandman and Reeve, 2006; Zlatanova, 1997). But the (H3-H4)-exclusive chromatin fiber will likely exhibit a strong bend. Comparison of the H3-H4 octasome with the archaeal nucleosome-like particle, known as the archaeasome, indicates a much wider DNA gyre separation (12.7 Å). This is because the inter-disk space of the H3-H4 octasome is occupied by the two extended, unstructured H3 N-termini, whereas in the archaeasome, the disks are held together by stacking interaction contributed in part by the inter-disk L1-L1’ contact between the dimers at positions N and N+3 (Mattiroli et al., 2017a). Interestingly, a recent study that combined cryo-EM and molecular simulation analyses indicates that archaeal chromatin is by no means a straight rod but is perhaps more slinky-like (Bowerman et al., 2021). In fact, the archaeasome disks can open, like a clam, with a 90° angle (Bowerman et al., 2021). Therefore, eukaryotic chromatin may share more similarity with archaea than previously thought.

Altogether, the structural insights of the H3-H4 octasome provided in this study underscore how eukaryotes may utilize alternative histone arrangements to modulate chromatin structure and dynamics. The next major challenge is to understand how H3-H4 octasomes interact with nuclear factors to modulate genomic functions.

## Materials and methods

### Purification of DNA fragments

The 145 bp DNA fragment derived from the Widom 601 sequence (ATCAGAATCCCGGTGCCGAGGCCGCTCAATTGGTCGTAGACAGCTCTAGCACCGCTTAAACGCACGTACGCGCTGTCCCCCGCGTTTTAACCGCCAAGGGGATTACTCCCTAGTCTCCAGGCACGTGTCAGATATATACATCGAT) was prepared as previously described (Dyer et al., 2003). Tandem repeats of the DNA fragment were inserted into the pGEM-T Easy vector (Promega). The plasmid was amplified in *Escherichia coli* cells and purified. The DNA fragment was cleaved by digestion with *Eco*RV (Takara), and the vector DNA region was removed by precipitation with polyethylene glycol (PEG) 6,000. The DNA fragment was further purified by TSKgel DEAE-5PM (TOSOH) column chromatography.

### Preparation of human histones, histone complexes, and nucleosomes

The human histones (H2A, H2B, H3.1, H3.1_ R49C_C96S_C110A, and H4) were expressed and purified by the method described previously (Tachiwana et al., 2010). The histone octamer and the H3-H4 tetramer were refolded by dialysis and purified by gel filtration chromatography on a HiLoad 16/600 Superdex 200 pg (Cytiva) column. The H3-H4 octasome and tetrasome were reconstituted with the H3-H4 tetramer, together with the 145 base-pair DNA, by the salt dialysis method (Tachiwana et al., 2010). The purified DNA was mixed with the H3-H4 tetramer in a DNA:histone molar ratio of 1:2.2 (for H3-H4 octasome) or 1:1 (for tetrasome) in 2 M KCl buffer, and the mixtures were dialyzed against buffer containing 10 mM Tris-HCl (pH 7.5), 2 M KCl, 1 mM EDTA, and 1 mM dithiothreitol (DTT). The KCl concentration was gradually reduced to 0.25 M by exchanging the buffer with a peristaltic pump. The reconstituted H3-H4 octasome (but not the tetrasome) was incubated at 55°C for 2 h and then purified by 6% non-denaturing PAGE using a Prep Cell apparatus (Bio-Rad).

### Micrococcal nuclease treatment assay

Nucleosomes with the 145 base-pair DNA (200 ng each) were mixed with 0.01 unit/μL micrococcal nuclease (MNase; Takara) and incubated at 37°C for 0, 3, 6, 18, and 36 min, in buffer containing 50 mM Tris-HCl (pH 7.5), 2.5 mM CaCl_2_, 1.9 mM dithiothreitol, and 50 mM NaCl. After the incubation, the reaction was stopped by adding deproteinization solution [20 mM Tris-HCl (pH 7.5), 20 mM EDTA, 0.5 mg/ml proteinase K, and 0.25% SDS]. The resulting DNA fragments were subjected to phenol/chloroform extraction and ethanol precipitation, and then analyzed by 10% native PAGE in 0.5x TBE buffer [45 mM Tris base, 45 mM boric acid and 1 mM EDTA].

### Native electrospray ionization (ESI) mass spectrometry

ESI mass spectra were acquired with a Triwave SYNAPT G2 HDMS mass spectrometer (Waters, Milford, MA) with a nanoESI source (Azegami et al., 2013; Giles et al., 2011; Saikusa et al., 2018; Zhong et al., 2011). For native ESI mass spectrometry, the H3-H4 octasome and tetrasome were dialyzed against 50 mM ammonium acetate (NH_4_OAc), and their concentrations were adjusted to 2 μM. The samples were deposited in a nanospray tip prepared in-house, and then placed in the nanoESI source. To observe the ions of the samples, the backing pressure was increased up to ~5 mbar by closing the SpeediValve. The following parameters were used for analysis: 0.70−1.0 kV capillary voltage, 20 V sampling cone voltage, 4 V trap collision energy, and 3.0 mL/min trap Ar-gas flow rate. The mass spectra were acquired for m/z 2,000–14,000, and calibrated with (CsI)nCs^+^ ions. The MassLynx version 4.1 software (Waters) was used for data processing and peak integration.

### Cryo-EM sample preparation and data collection

To stabilize the purified H3-H4 octasome, the gradient fixation method (GraFix) was performed using sucrose gradient centrifugation (Kastner et al., 2008). The gradient solution was prepared using a Gradient Master (SKB) with low-sucrose buffer [20 mM HEPES-KOH (pH 7.5), 1 mM DTT, and 10% (w/v) sucrose] and high-sucrose buffer [20 mM HEPES-KOH (pH 7.5), 1 mM DTT, 25% (w/v) sucrose, and 3% paraformaldehyde]. Centrifugation was performed for 16 h at 27,000 rpm at 4°C, using an SW41 rotor (Beckman Coulter). The fractions containing the H3-H4 octasome were buffer-exchanged to 20 mM Tris-HCl (pH 7.5) buffer containing 1 mM DTT, using Micro Bio-Spin Columns (Bio-Rad). For cryo-EM specimen preparation, a 2 μL aliquot of the H3-H4 octasome (0.63 mg/mL) was applied to a glow-discharged Quantifoil R1.2/1.3 200-mesh Cu grid and blotted for 8 seconds under 100% humidity at 12°C in a Vitrobot Mark IV (Thermo Fisher Scientific, USA). The grids with the H3-H4 octasome were immediately plunged into liquid ethane. Cryo-EM data collection of the H3-H4 octasome was performed by the SerialEM auto acquisition software (Mastronarde, 2005) on a Krios G3i cryo-electron microscope (Thermo Fisher Scientific, USA) operated at 300 kV at a pixel size of 1.05 Å and a defocus range from −1.25 to −2.5 μm. Images of the H3-H4 octasome were recorded with 6 second exposure times on an energy-filtered K3 direct electron detector (Gatan, USA) in the electron counting mode with a slit width of 25 eV, retaining a total of 40 frames with an overall dose of ~63 electron/Å^2^.

### Cryo-EM image processing

In total, 5,517 movies of the H3-H4 octasome were aligned by MOTIONCOR2 (Zheng et al., 2017) with dose weighting. The contrast transfer function (CTF) parameters for each micrograph were estimated by CTFFIND4 (Rohou and Grigorieff, 2015). RELION 3.0 (Zivanov et al., 2018) was used for the following image processing of the H3-H4 octasome. From 3,922 micrographs, 1,427,558 particles were picked automatically and subjected to reference-free 2D classification. Subsequently, the selected 990,623 particles were used for the 3D classification. The two best classes from the 3D classification were selected based on the resolution and subjected to the 3D refinement, followed by Bayesian polishing and two rounds of CTF refinement. The final resolutions of the refined maps of the closed and open conformations of the H3-H4 octasome, based on the gold standard Fourier Shell Correlation (FSC) with the 0.143 criterion (Scheres, 2016), were 3.6 Å and 3.9 Å, respectively. The final maps of the H3-H4 octasome were normalized with MAPMAN (Kleywegt et al., 2004) and visualized with UCSF Chimera (Pettersen et al., 2004). The details of the processing statistics for the H3-H4 octasome are presented in Table 1.

### Model building

The structural models of the closed and open forms for the H3-H4 octasome were built from the H3-H4 tetramer and its proximal DNA fragment in the crystal structure of the nucleosome containing *Xenopus laevis* histones and a 145 base-pair Widom 601 DNA (PDB: 3LZ0), which represented roughly half of the symmetric structures (Vasudevan et al., 2010). The amino acid residues of the histones were adjusted to those of human histones. The model coordinates were refined automatically with phenix.real_space_refine (Adams et al., 2010) and manually using Coot (Emsley et al., 2010). The DNA sequence of the H3-H4 octasome was estimated based on the MNase digestion, followed by the restriction enzyme analysis of the H3-H4 octasome. All structure figures were prepared using UCSF Chimera and PyMOL (Schrödinger; http://www.pymol.org).

### Yeast plasmids

Yeast histone expression vectors were derived from the *HHT1-HHF1 URA3 CEN ARS* plasmid pMS329 (gift of Mitchell Smith) and the *HHT2-HHF2 TRP1 CEN ARS* plasmid pWZ414-F12 (obtained from Rolf Sternglanz) (Megee et al., 1990; Zhang, 1998). The *HHT1(R49C)* vector (pEL649) and the *HHT2(R49C)* (pEL629) vector were constructed by digesting pMS329 and pWZ414-F12 respectively with ClaI/AgeI and with *Blp*I/*Bam*HI to remove a segment of their H3 genes followed by recombination with synthetic DNA fragments (Twist Bioscience) containing the *R49C* mutation by the Gibson assembly protocol (New England Biolabs). Similarly, the 2xV5-*HHT1* (pEL656) and 2xV5-*HHT1(R49C)* (pEL650) plasmids were generated by digesting pMS329 and pEL649, respectively, with AgeI/PmeI followed by recombination with a synthetic fragment containing the N-terminal 2xV5 tag. The integrity of all plasmids was confirmed by Sanger sequencing (Genewiz).

### Yeast strains

The histone knockout strain YYY67 (gift of Rolf Sternglanz) was supplemented with pMS329 (Megee et al., 1990; Yu et al., 2011). The *HHT2(R49C)* allele and the *HHT2* control were introduced into YYY67 along with *HHF2* as *TRP1 CEN ARS* plasmids. Transformants were selected on synthetic complete (SC) media lacking uracil and tryptophan and then seeded onto complete synthetic media (CSM) supplemented with 0.1% (w/v) 5-fluoroorotic acid (5-FOA) to select against the *URA3* plasmid(Megee et al., 1990). Survivors of the 5-FOA selection represent the wild-type *HHT2* strain (yEL699) and the mutant *HHT2(R49C)* strain (yEL705) used in Figure 4C. For the hybrid mutant in figure supplement S3, *HHT1(R49C)* and *HHT1* with or without an N-terminal 2xV5 tag were introduced into yEL699 and yEL705 on *URA3 CEN ARS* plasmids. When generating the *HHT2(R49C)* yEL705 strain, a strong growth defect was observed on the 5-FOA media, but the defect was partially alleviated when the cells were grown anaerobically using the anaerobe pouch system (BD Biosciences, cat# B260683). The growth defect of *HHT2(R49C)* was less severe in media without 5-FOA, and thus the cells were grown aerobically for VivosX.

### VivosX

VivosX was performed as reported previously (Mohan et al., 2018). Briefly, yeast cells expressing *HHT2-HHF2*(yEL699) or *HHT2(R49C)-HHF2*(yEL705) were cultured at 30°C to an optical density (at 600 nm) of ~0.5 before treatment with 180 μM 4-DPS (in DMSO) or with an equivalent volume of DMSO for 20 min at 30°C. The cells (from 5 mL of the culture) were fixed with 20% trichloroacetic acid (TCA), pelleted by centrifugation, and homogenized in 20% TCA by Zirconia bead beating using a FastPrep-24 machine. The precipitates were washed with acetone and extracted with 200 μL TUNES-GN buffer [100 mM Tris-HCl (pH 7.2), 6 M urea, 10 mM EDTA, 1% SDS, 0.4 M NaCl, 10% glycerol, and 50 mM N-ethylmaleimide (NEM)] at 30°C for 1 h with vortexing. SDS-PAGE was performed under non-reducing or reducing conditions as previously described (Mohan et al., 2018). H3 and V5 western analyses were performed with an anti-H3 antibody (gift of Carl Wu) and an anti-V5 antibody (Thermo 46-0705) at 1:2,000 dilution. The anti-H3 antibody used in figure supplement 3C was affinity purified. Secondary antibodies conjugated to horseradish peroxidase were used at 1:5,000 dilution. Western signals were developed with the ECL Prime reagent (GE Life Sciences, RPN2232) and imaged with an LAS-4010 CCD camera system (GE Life Sciences).

### In vitro disulfide crosslinking analysis

Nucleosomes and H3-H4 octasomes were reconstituted with human histones containing the C96S, C110A, and R49C substitutions on H3.1 in 20 mM Tris-HCl (pH 7.0). The nucleoproteins were mixed with 4-DPS (or without) to a final concentration of 0.8 μM and incubated at 25°C for 30 min. For reducing SDS-PAGE, the reaction aliquots were mixed (1:1) with a sample buffer containing 100 mM Tris-HCl (pH 6.8), 4% SDS, 20% glycerol, 0.2% BPB, and 200 mM 2-mercaptoethanol. The samples (280 ng each) were heated at 95°C for 5 min before they were analyzed by 16 % polyacrylamide SDS-PAGE and Coomassie Brilliant Blue staining. For non-reducing SDS-PAGE, the reaction aliquots were mixed with a similar sample buffer but without bME and analyzed by SDS-PAGE without prior heating.

## Acknowledgments

We are grateful to the members of the Kurumizaka laboratory, particularly Keiko Gotanda, Lumi Negishi, Yukari Iikura and Yas Takeda, for their assistance and to the members of the Luk lab for critical reading of the manuscript. We thank Akihisa Tsutsumi, of Masahide Kikkawa’s laboratory (The University of Tokyo), for cryo-EM data collection.

## Author contributions

K.N. prepared the nucleosomes and carried out biochemical analyses. K.N. and Y.T. performed cryo-EM analyses. K.S. and S.A. performed ESI-MS experiments. P.L. and E.L. prepared yeast strains and performed VivosX experiments. H.K. and E.L. designed and supervised the research. K.N., E.L. and H.K. interpreted the data and wrote the manuscript. All of the authors discussed the results and commented on the manuscript.

## Additional files

### Supplementary files

Supplementary file: figure supplements 1-3, Tables 1-3

## Data and materials availability

The cryo-EM reconstructions and atomic models of the H3-H4 octasome have been deposited in the Electron Microscopy Data Bank and the Protein Data Bank (PDB) under the following accession codes: EMD-XXXXX and YYYY for closed form, and EMD-ZZZZZ and WWWW for open form.

