## supplemental data for "Cryo-electron microscopy structure of the H3-H4 octasome without histones H2A and H2B"

**This PDF file includes:**

figure supplements 1-3

Tables 1-3

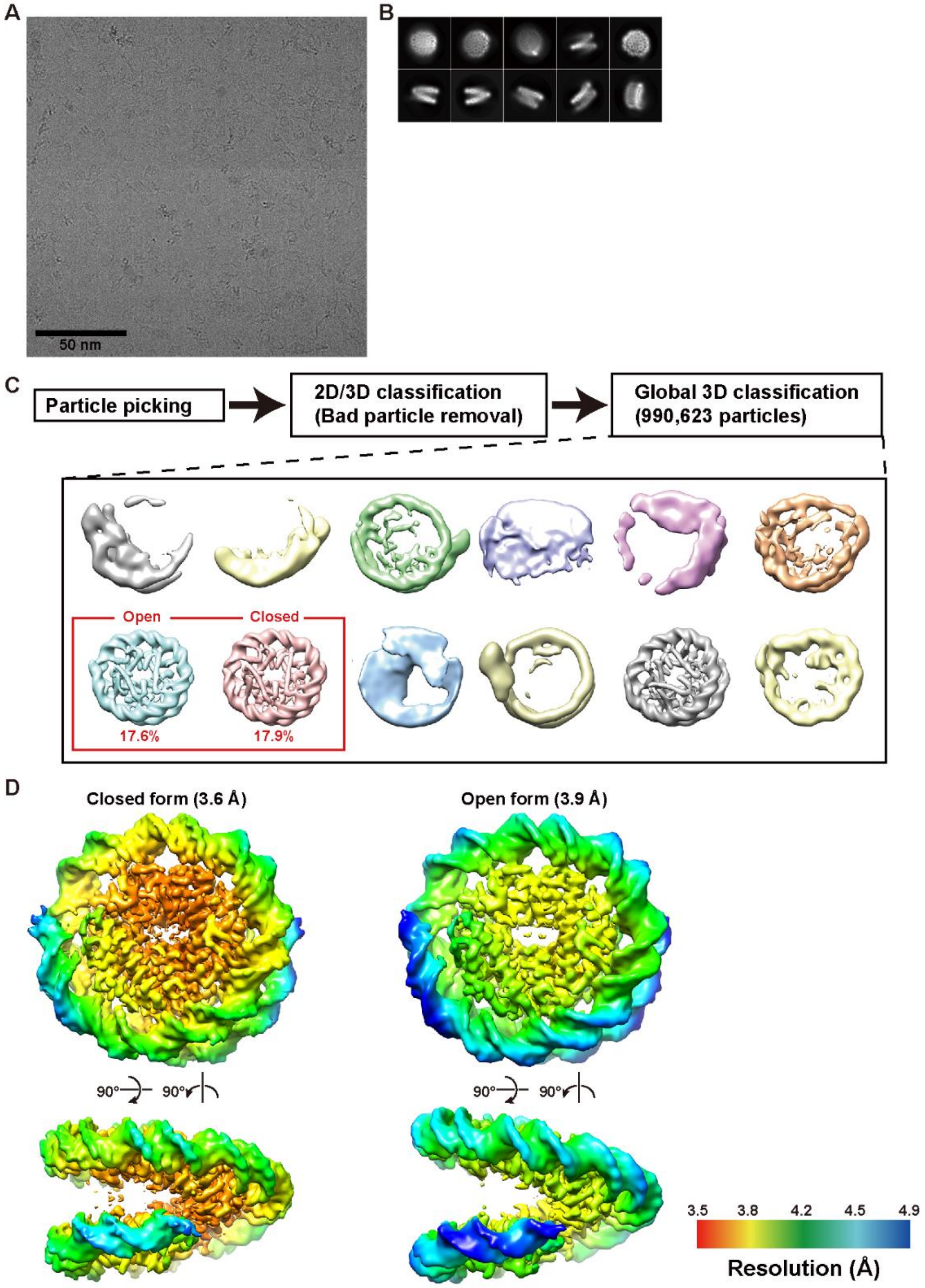


**figure supplement 1. Cryo-EM data collection and initial image processing**

(A) Representative micrograph of the cryo-EM dataset. (B) Representative 2D class averages from the reference-free 2D classification calculated after removing bad particles. (C) Workflow of the initial stage of the image processing. Box size is 18.9 nm. (D) Local resolution maps for the closed (left panel) and open (right panel) forms of the H3-H4 octasome. Local resolutions were estimated using the RELION postprocess.

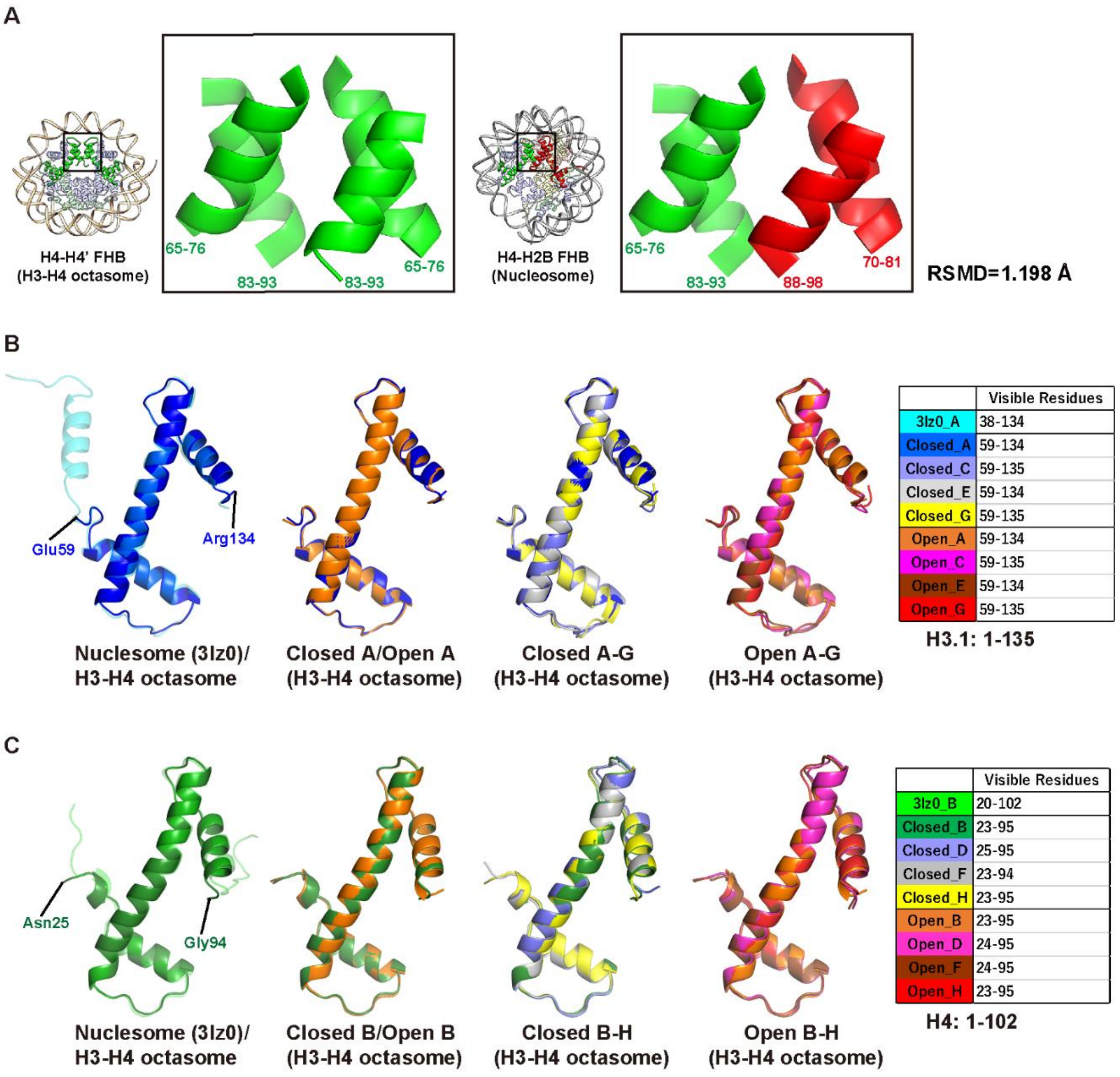


**figure supplement 2. Structural comparison of histones in H3-H4 octasome and nucleosome**

(A) Structural comparison of the H4-H4' FHB in the H3-H4 octasome (left panel) and the H4-H2B FHB in the nucleosome (right panel). These FHB structures are superimposable with a root mean square deviation (RMSD) of 1.198 Å of the backbone. (B) Structural comparison of H3.1 in the H3-H4 octasome and the nucleosome. (C) Structural comparison of the H4 in the H3-H4 octasome and the nucleosome.

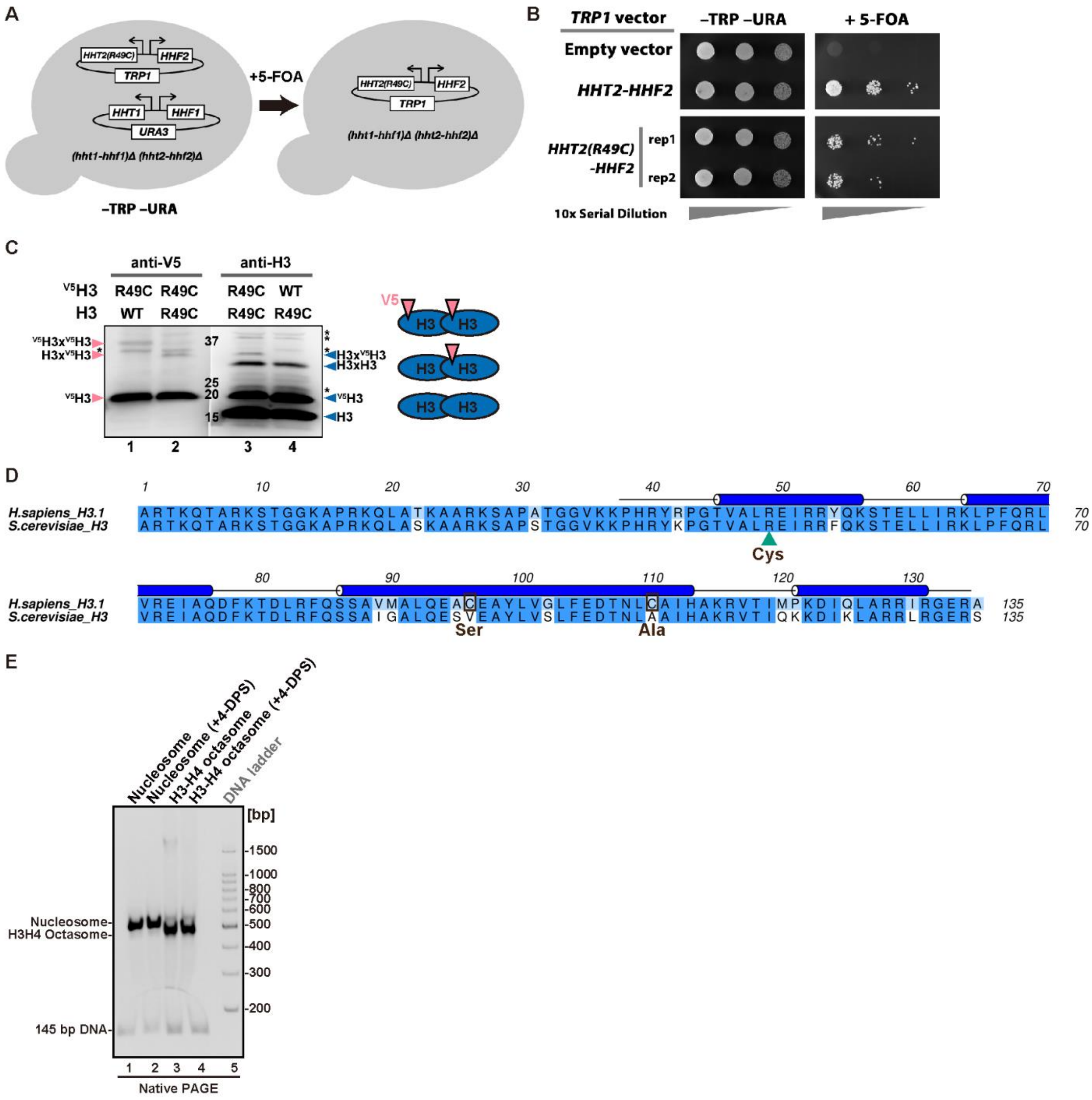


**figure supplement 3. Genetic and biochemical analyses of the H3-H4 octasome containing the H3 R49C mutant**

(A) The strategy used to verify the functionality of the *HHT2(R49C)* gene (Megee et al., 1990). (B) A plate test showing the growth of yeast expressing *HHT2(R49C)* or wild-type *HHT2* as the sole source of H3. The cells were grown at 30°C under anaerobic conditions to alleviate the growth defect of *HHT2(R49C)* in 5-FOA. (C) VivosX analysis of V5-tagged H3 (*HHT1*) and untagged H3 (*HHT2*) with R49C or without (WT) under non-reducing SDS-PAGE conditions. Samples 1-4 were analyzed on an 8-16% polyacrylamide gel and transferred to a PVDF membrane. The membrane was cut along the molecular weight marker lane (between samples 2 and 3). The left half of the membrane was probed with an anti-V5 antibody, the right half with an affinity-purified anti-H3 antibody. The V5 and H3 western images were re-aligned using the marker lane. Asterisks indicate non-specific bands. Blue arrowheads highlight the H3 species detected by anti-H3. Pink arrowheads highlight the V5-containing species. Note that *HHT2* is expressed at a higher level than *HHT1*. As a result, the H3-H3' crosslinked species is biased against the $^{V5}$H3(Hht1)-linked species on the anti-H3 blot. The $^{V5}$H3x$^{V5}$H3 adduct, which is indicated by the anti-V5 blot, is almost undetectable by the anti-H3 western. (D) Sequence alignment analysis between the human H3.1 and yeast H3 proteins. The H3.1 mutations used in the *in vitro* crosslinking experiment are indicated with a green triangle and black squares. (E) Reconstituted nucleosomes and H3-H4 octamers (400 ng each) containing the H3.1 C96S C110A R49C mutant with or without 4-DPS were analyzed by 6% polyacrylamide native PAGE with ethidium bromide (EtBr) staining.

**Table 1: Cryo-EM processing statistics**

| Sample (H3-H4 octasome) | Closed form (EMD-XXXXX) (PDB: YYYY) | Open form (EMD-ZZZZZ) (PDB: WWWW) |
|---|---|---|
| **Data collection** | | |
| Electron microscope | Krios G3i | Krios G3i |
| Camera | K3 | K3 |
| Pixel size (Å/pix) | 1.05 | 1.05 |
| Defocus range (µm) | -1.25 to -2.5 | -1.25 to -2.5 |
| Exposure time (second) | 6 | 6 |
| Total dose (e/Å$^2$) | 63 | 63 |
| Movie frames (no.) | 40 | 40 |
| Total micrographs (no.) | 5,517 | 5,517 |
| **Reconstruction** | | |
| Software | Relion 3.0 | Relion 3.0 |
| Particles for 2D classification | 1,427,558 | 1,427,558 |
| Particles for 3D classification | 990,623 | 990,623 |
| Particles in the final map (no.) | 186,253 | 177,542 |
| Symmetry | C2 | C2 |
| Final resolution (Å) | 3.6 | 3.9 |
| FSC threshold | 0.143 | 0.143 |
| Map sharpening B factor (Å$^2$) | -37.53 | -35.87 |
| **Model building** | | |
| Software | Coot | Coot |
| **Refinement** | | |
| Software | Phenix | Phenix |
| **Model composition** | | |
| Protein | 980 | 980 |
| Nucleotide | 290 | 290 |
| **Validation** | | |
| MolProbity score | 1.52 | 1.49 |
| Clash score | 9.9 | 9.15 |
| R.m.s. deviations | | |
| Bond lengths (Å) | 0.004 | 0.004 |
| Bond angles (˚) | 0.783 | 0.750 |
| **Ramachandran plot** | | |
| Favored (%) | 98.62 | 98.62 |
| Allowed (%) | 1.38 | 1.38 |
| Outliers (%) | 0 | 0 |

### Table 2: Yeast Strains

| Strain | Genotype | Source |
|---|---|---|
| **YYY67** | MATa leu2Δ1 his3Δ200 ura3-52 trp1Δ63 lys2-128δ (hht1-hhf1)Δ::LEU2 (hht2-hhf2)Δ::HIS3 Ty912Δ35-lacZ::his4 <pMS329 (URA3 SUP11 CEN4 ARS4 HHT1-HHF1)> | (Yu et al., 2011) |
| **yEL690** | MATa leu2Δ1 his3Δ200 ura3-52 trp1Δ63 lys2-128δ (hht1-hhf1)Δ::LEU2 (hht2-hhf2)Δ::HIS3 Ty912Δ35-lacZ::his4 <pMS329 (URA3 SUP11 CEN4 ARS4 HHT1-HHF1)> <pRS414 (TRP1 CEN6 ARS4)> | This study |
| **yEL691** | MATa leu2Δ1 his3Δ200 ura3-52 trp1Δ63 lys2-128δ (hht1-hhf1)Δ::LEU2 (hht2-hhf2)Δ::HIS3 Ty912Δ35-lacZ::his4 <pMS329 (URA3 SUP11 CEN4 ARS4 HHT1-HHF1)> <pWZ414-F12 (TRP1 CEN6 ARS4 HHT2-HHF2)> | This study |
| **yEL698** | MATa leu2Δ1 his3Δ200 ura3-52 trp1Δ63 lys2-128δ (hht1-hhf1)Δ::LEU2 (hht2-hhf2)Δ::HIS3 Ty912Δ35-lacZ::his4 <pMS329 (URA3 SUP11 CEN4 ARS4 HHT1-HHF1)> <pEL629 (TRP1 CEN6 ARS4 HHT2(R49C)-HHF2)> | This study |
| **yEL699** | MATa leu2Δ1 his3Δ200 ura3-52 trp1Δ63 lys2-128δ (hht1-hhf1)Δ::LEU2 (hht2-hhf2)Δ::HIS3 Ty912Δ35-lacZ::his4 <pWZ414-F12 (TRP1 CEN6 ARS4 HHT2-HHF2)> | This study |
| **yEL705** | MATa leu2Δ1 his3Δ200 ura3-52 trp1Δ63 lys2-128δ (hht1-hhf1)Δ::LEU2 (hht2-hhf2)Δ::HIS3 Ty912Δ35-lacZ::his4 <pEL629 (TRP1 CEN6 ARS4 HHT2(R49C)-HHF2)> | This study |
| **yEL728** | MAT a leu2Δ1 his3Δ200 ura3-52 trp1Δ63 lys2-128δ (hht1-hhf1)Δ::LEU2 (hht2-hhf2)Δ::HIS3 Ty912Δ35-lacZ::his4 <pEL650 (URA3 SUP11 CEN4 ARS4 2xV5-hht1(R49C)-HHF1)> <pWZ414-F12 (TRP1 CEN6 ARS4 HHT2-HHF2)> | This study |
| **yEL729** | MAT a leu2Δ1 his3Δ200 ura3-52 trp1Δ63 lys2-128δ (hht1-hhf1)Δ::LEU2 (hht2-hhf2)Δ::HIS3 Ty912Δ35-lacZ::his4 <pEL656 (URA3 SUP11 CEN4 ARS4 2xV5-HHT1-HHF1)> <pEL629 (TRP1 CEN6 ARS4 HHT2(R49C)-HHF2)> | This study |
| **yEL730** | MAT a leu2Δ1 his3Δ200 ura3-52 trp1Δ63 lys2-128δ (hht1-hhf1)Δ::LEU2 (hht2-hhf2)Δ::HIS3 Ty912Δ35-lacZ::his4 <pEL650 (URA3 SUP11 CEN4 ARS4 2xV5-hht1(R49C)-HHF1> <pEL629 (TRP1 CEN6 ARS4 HHT2(R49C)-HHF2)> | This study |

**Table 3: Yeast Plasmids**

| ***Plasmid*** | ***Description*** | ***Source*** |
|---|---|---|
| pMS329 | *URA3 SUP11 CEN4 HHT1-HHF1* | (Megee et al., 1990) |
| pRS414 | *TRP1 CEN6 ARS4* | (Sikorski and Hieter, 1989) |
| pWZ414-F12 | *TRP1 CEN6 ARS4 HHT2-HHF2* | (Zhang, 1998) |
| pEL629 | *TRP1 CEN6 ARS4 HHT2(R49C)-HHF2* | This study |
| pEL649 | *URA3 SUP11 CEN4 HHT1(R49C)-HHF1* | This study |
| pEL650 | *URA3 SUP11 CEN4 2xV5-HHT1(R49C)-HHF1* | This study |
| pEL656 | *URA3 SUP11 CEN4 2xV5-HHT1-HHF1* | This study |
